# Simultaneous Dissection of Grain Carotenoid Levels and Kernel Color in Biparental Maize Populations with Yellow-to-Orange Grain

**DOI:** 10.1101/2021.09.01.458275

**Authors:** Mary-Francis LaPorte, Mishi Vachev, Matthew Fenn, Christine Diepenbrock

## Abstract

Maize enriched in provitamin A carotenoids could be key in combatting vitamin A deficiency in human populations relying on maize as a food staple. Consumer studies indicate that orange maize may be regarded as novel and preferred. This study identifies genes of relevance for grain carotenoid concentrations and kernel color, through simultaneous dissection of these traits in 10 families of the U.S. maize nested association mapping population that have yellow to orange grain. Quantitative trait loci (QTL) were identified via joint-linkage analysis, with phenotypic variation explained for individual kernel color QTL ranging from 2.4 to 17.5%. These QTL were cross-analyzed with significant marker-trait associations in a genome-wide association study that utilized ∼27 million variants. Nine genes were identified: four encoding activities upstream of the core carotenoid pathway, one at the pathway branchpoint, three within the α- or β-pathway branches, and one encoding a carotenoid cleavage dioxygenase. Of these, three exhibited significant pleiotropy between kernel color and one or more carotenoid traits. Kernel color exhibited moderate positive correlations with β-branch and total carotenoids and negligible correlations with α-branch carotenoids. These findings can be leveraged to simultaneously achieve desirable kernel color phenotypes and increase concentrations of provitamin A and other priority carotenoids.

## INTRODUCTION

Kernel color and grain carotenoid profiles are valuable traits that directly impact consumer preference and crop nutritional quality in maize (*Zea mays* ssp. *mays* L.). Maize is a cereal crop ranking globally among the most important sources of daily calories, and is estimated to provide 38% of the food supply in Africa (Prasanna et al. 2020; Chandler et al. 2013); as such, maize has been a key crop for biofortification efforts. Some carotenoid compounds are pigments, and overall abundance of maize grain carotenoids has been found to exhibit weak positive correlations with kernel color in 228 diverse inbreds (R^2^ = 0.119; Harjes et al. 2008). However, genetic and phenotypic relationships of several priority carotenoid traits in maize grain and kernel color have not yet been dissected in tandem. Achieving a greater understanding of these relationships through simultaneous examination of these traits in the same experimental framework (i.e., populations and environments) could expedite efforts to select for deep orange kernel color and improved carotenoid profiles in tandem, for the development of multi-value added products.

As humans are unable to endogenously synthesize vitamin A, dietary intake of provitamin A is crucial for proper immune system development and healthy vision (Tanumihardjo et al. 2016). Three carotenoid compounds have provitamin A activity. β-carotene provides two units of retinol (active vitamin A) upon oxidative cleavage in human and animal systems. β-cryptoxanthin provides one unit of retinol but recent evidence has suggested its greater bioavailability compared to β-carotene (Prasanna et al. 2020), such that the two compounds are now considered equivalent in breeding efforts. Finally, α-carotene provides one unit of retinol. In the case of young children going through pre-adolescent development, sufficient vitamin A intake is especially important; deficiency can result in impaired immunity and stunted growth, blindness, and ultimately death (Underwood and Arthur 1996). Over 95% of deaths in children due to vitamin A deficiency occur in sub-Saharan Africa or south Asia (Stevens et al. 2015). Furthermore, non-fatal vitamin A deficiencies contribute to permanent corneal scarring and/or night blindness, resulting in lifelong effects (Stevens et al. 2015). Alleviating vitamin A deficiency in both children and adults must continue to be a high-priority target for improving the human condition worldwide.

The improvement of crop nutritional quality through plant breeding and/or agronomic strategies, termed biofortification, could serve as a viable complement or alternative to other approaches to address nutritional deficiencies in certain subregional contexts (Welch and Graham 1999). The success of approaches such as crop and dietary diversification, processing-stage fortification, or supplementation (Mora 2003; Ross 2002; West 2000; West et al. 2002) can be limited by requirements for crop growth or by difficulties arising in transportation and infrastructure, among other factors (Graham et al. 2001; Gadaga et al. 2009). Biofortification of maize with provitamin A has been found to be a cost-effective and sustainable approach (Bouis and Welch 2010), and extensive natural variation has been observed for grain carotenoid traits among diverse maize accessions (Owens et al. 2014), which can be leveraged in breeding efforts.

Carotenoids are primarily localized in the endosperm of maize grain and are biosynthesized in the plastids. The methyl-D-erythritol-4-phosphate (MEP) pathway produces isopentenyl pyrophosphate (IPP), a precursor from which carotenoids and other plastidic isoprenoid compounds are derived (Cordoba et al. 2011). Flux through the core carotenoid biosynthetic pathway commences with the biosynthesis of phytoene from geranylgeranyl diphosphate (GGDP), a step catalyzed by phytoene synthase (Yuan et al. 2015). Subsequent desaturations convert the backbone of phytoene into a light-absorbing chromophore composed of repeating conjugated double bonds (Bartley and Scolnik 1995), and sequential desaturation and isomerization reactions produce all-*trans* lycopene; all-*trans* is the predominant isomer among lycopene and most other carotenoids. The addition of a β- and ε-ring or, alternatively, two β- rings to each terminal carbon of lycopene yield α- and β-carotene, respectively, and further hydroxylation will generate xanthophylls such as lutein or zeaxanthin (Khoo et al. 2011; Yuan et al. 2011).

Two carotenoid biosynthetic genes in particular have been utilized in maize provitamin A biofortification efforts thus far: *lycopene epsilon cyclase (lcyE)*, which adds the ε-ring to lycopene and represents a key branchpoint of the carotenoid pathway, and *beta carotene hydroxylase 1* (*crtRB1*), which converts β-carotene to β-cryptoxanthin and then zeaxanthin (Pixley et al. 2013; Prasanna et al. 2020). Using these genes of interest, researchers affiliated with the International Maize and Wheat Improvement Center (CIMMYT), the International Institute of Tropical Agriculture (IITA), HarvestPlus, and partners have developed maize varieties that accumulate higher levels of provitamin A carotenoids in the grain and are regionally adapted to different growing conditions and endemic stressors (Pixley et al. 2013; Menkir et al. 2017; Prasanna et al. 2020). Lutein and zeaxanthin, while not having provitamin A activity, are also priority carotenoids for human health given their important—including protective—roles as major constituents of the macular pigment of the eye (Beatty et al. 1999, Krinsky et al. 2003, Bernstein and Arunkumar 2021). Increased dietary intake of these compounds has been associated with lower risk of age-related macular degeneration (AMD; Abdel-Aal et al. 2013, Bernstein and Arunkumar 2021). AMD is of global significance as a common cause of vision loss and blindness in adults. A total of 170 million adults were estimated to be affected in 2014, with projected increases in prevalence by 2040 (Wong et al. 2014).

While biofortification of maize with provitamin A and other priority carotenoids (such as lutein and zeaxanthin) is a promising opportunity to address deficiency and reduce global disease burdens, it has also been critical to consider consumer preference with regards to kernel color. In parts of eastern and southern Africa, white maize is preferred for human consumption, whereas yellow maize is dispreferred, including due to historical issues associated with long storage periods for shipments of yellow maize (De Groote and Kimenju 2008; Pillay et al. 2011; Tschirley and Santos 1994; Muzhingi et al. 2008). One solution, given this dispreference for yellow kernel color, is to develop orange-grained maize varieties. Stevens and Winter-Nelson (2008) found that consumers surveyed in Mozambique did not have an aversion to orange biofortified maize, and on average rated its aroma more favorably than the alternative white maize. This study suggested that consumer preference for the orange biofortified maize may have been due to educational presentations in which the surveyors explained the maize’s nutritional benefits, and that consumers may be attracted to the product for its ability to alleviate vitamin A deficiency.

The relationship between kernel color and grain carotenoid concentrations is not sufficiently consistent for orange kernel color to be selected upon for the improvement of concentrations of provitamin A and other priority carotenoids (Pfeiffer and McClafferty 2007; Harjes et al. 2008). This is due in part to the identity and spectral properties of the carotenoids that accumulate in maize endosperm and certain aspects of their relative abundance—namely, lutein and zeaxanthin tend to be the most abundant carotenoids in maize grain (Owens et al. 2014) and are themselves pigmented, which could tend to mask impacts of other carotenoid compounds on kernel color. However, the nature and extent of relationships between the overall abundance vs. relative abundance (i.e., composition) of several carotenoid compounds with kernel color has not yet been dissected in the same quantitative genetics-enabling experimental framework, and will be tested herein. Additionally, genes previously identified for quantitative components of kernel color in 1,651 diverse maize inbreds (Owens et al. 2019) have been implicated in both overall carotenoid abundance and composition in other experimental frameworks (Owens et al. 2014, Diepenbrock et al. 2021), which could complicate the achievement of maximized tandem gains for provitamin A (among other priority) carotenoids and kernel color. This study aims to simultaneously dissect the genetic basis of—and genetic and phenotypic relationships among—eight grain carotenoid traits in tandem with visually scored kernel color, in 10 families of the U.S. maize nested association mapping (NAM) panel having yellow to orange grain color, to identify and characterize potential targets for the accelerated development of orange, carotenoid-dense maize.

## MATERIALS AND METHODS

### Plant Materials and Trait Quantification

The 10 families analyzed in this study are part of the 25-family U.S. maize nested association mapping (NAM) panel, which has been previously described (Yu et al. 2008; Buckler et al. 2009; McMullen et al. 2009). The experimental design for field evaluations of the NAM panel, conducted in the summers of 2009 and 2010 at the Purdue University Agronomy Center for Research and Education (West Lafayette, IN) using standard agronomic practices, was as described in Chandler et al. (2013). Briefly, the 2009 environment used two fields to grow the entire single replicate of the U.S. maize NAM panel, whereas the 2010 environment used one field to grow the entire single replicate. A sets design was used in each environment, in which each family was planted in a population block (or set). The set for each NAM family was planted in an augmented incomplete block □-lattice design with block size 10 × 20, with B73 and the alternate parent of that family used as the checks within each block. The 281-line Goodman-Buckler maize diversity panel was additionally planted in each environment, in an augmented incomplete block □-lattice design with block size 14 × 20 and with B73 and Mo17 as the checks within each block. For the analyses described herein, we utilized data that was collected on samples from population blocks corresponding to the 10 families of the U.S. maize NAM population that are of interest in this study, with B73 as the common parent and the following as alternate parents: B97, CML228, CML52, Hp301, Ki11, Ki3, NC350, NC358, Oh7B, and Tx303. Chandler et al. (2013) had identified these 10 families to be segregating for yellow to orange kernel color. A single-row plot containing approximately 10 plants of a single inbred (with plot length of 3.05 m) was the individual experimental unit for this study. Self-pollination was conducted by hand for at least four plants per plot. Kernel samples from these self-pollinated ears were harvested, dried to 15% moisture, shelled, and bulked to form a representative sample (per plot) for kernel color and carotenoid phenotyping.

Ordinal scores for kernel color, on a scale of 1 (yellow) to 12 (deep orange), were as evaluated for these 10 families in Chandler et al. (2013). Briefly, ordinal scores were assessed on a bulk sample of 100+ kernels per plot with the embryo facing down. Within a single bulk sample, kernels were grouped by color (if multiple kernel colors were present), and each group was then assigned a score. Most bulk samples were uniform in color. For those with multiple color groups, the score recorded for that bulk sample (representing a single plot) was the average of the score assigned to each group. Each bulked sample was scored once, by the same person. Bulked samples in which kernels were discolored due to fungal and/or other opportunistic pathogens did not receive a color score. For bulked samples in which the pericarp was pigmented due to anthocyanins, or if kernel color was otherwise difficult to score without ambiguity, the pericarp of the kernels was removed prior to scoring. Carotenoid phenotypes (μg g^−1^ seed) were a subset of those analyzed in Diepenbrock et al. (2021), which were collected via high-performance liquid chromatography (HPLC) as described in that study, on ∼50 ground seeds per plot from the same field experiment. 20 μL of seed extract was injected for each sample, with 1 mg of β-apo-8′-carotenal used as an internal standard. External standards were used in five-point curves for quantification of carotenoid compounds at 450 nm, with a lutein curve used for lutein, zeinoxanthin, and α-carotene; a β-carotene curve for β-carotene and β-cryptoxanthin; and a zeaxanthin curve for zeaxanthin. Relative phytofluene levels (at 350 nm) were also estimated from the β-carotene curve.

### Best linear unbiased estimators

Using the phenotypic data from the 10 families, best linear unbiased estimators (BLUEs) were generated from the Chandler et al. (2013) kernel color phenotypes and Diepenbrock et al. (2021) carotenoid phenotypes using a process similar to that taken in Diepenbrock et al. (2021). Briefly, family, RIL within family, year, and field within year were fitted as random effects as a baseline model. The best random structure was then determined using the Bayesian Information Criterion (BIC; Schwarz 1978), to determine which of zero to five additional terms were optimally included in the final model. The additional terms tested were HPLC auto-sampler plate, set within field within year, block within set within field within year, family within year, and RIL within family within year. The best residual structure was again determined using BIC, conditional upon the best random structure. The residual structures tested included identity by year; autoregressive for range and identity for row, by field-in-year; identity for range and autoregressive for row, by field-in-year; and autoregressive (first-order, AR1 × AR1) for range and row, by field-in-year. Field-in-year was a new factor that combined the field used and the year, to enable fitting of a separate error structure for each of the three fields used in this experiment. The final model using the optimal random and residual structures was then fit, and outlying observations (for a given RIL or check genotype, trait, and field-in-year combination) were identified via difference in fits (DFFITs; Neter et al. 1996; Belsley et al. 2005) and were set to NA for that specific trait if exceeding a conservative threshold previously found to be appropriate for this experimental design (Hung et al. 2012). The final model was then fit again with RIL and RIL within family now specified as sparse fixed effects rather than random effects, for the generation of BLUEs. The BLUEs then underwent Box-Cox transformation using the optimal convenient lambda identified for each trait (Table S1). Untransformed and transformed BLUEs are reported in Table S2. The final model for each trait was also then fit with all terms specified as random effects except for the grand mean, to produce variance components to be used in the estimation of heritabilities on a line-mean basis. These heritabilities were estimated across the 10 families in this study (Hung et al. 2012), and the delta method was used to determine standard errors (Holland et al. 2003).

### Joint-linkage analysis

Joint-linkage (JL) analysis was conducted in TASSEL5 as described in Diepenbrock et al. (2021), using a 0.1 cM consensus genetic linkage map consisting of 14,772 markers. This map was generated by imputing SNP markers at 0.1 cM intervals, anchored on genotyping-by-sequencing data for the ∼4,900 RILs of the U.S. maize NAM population. The JL analysis was performed on transformed BLUEs of each trait, with the family term forced into the model first as a predictor variable. Each of the 14,772 markers nested within family were then tested for potential inclusion in the model as a predictor variable via joint stepwise regression. Entry thresholds were determined for each trait by conducting 1,000 permutations and selecting the *P-*value (from a partial F-test) corresponding to a Type I error rate of α = 0.05. Exit thresholds were set to equal twice the entry threshold, so that entry and exit of a given marker could not take place in the same step. One pair of multicollinear SNPs (defined as magnitude of Pearson’s correlation of SNP genotype state scores being greater than 0.8) was identified for lutein. The SNP having the lower sum of squares was removed from the JL model, and the model was fit again with re-scan in the vicinity of the remaining peak markers in the model. Namely, if another marker in the respective support interval now exhibited a larger sum of squares than the originally identified peak marker, it would be included in the model instead and the support interval re-calculated, until a local maximum in the sum of squares was identified. Once the final model was fit, the allelic effect estimates were calculated, nested within family, as described in Diepenbrock et al. (2017). Specifically, the final JL models were fitted using the lm() function from the lme4 package (Bates et al. 2015; R Core Team 2018) (Table S3). Phenotypic variation explained (PVE) was then calculated as in Li. et al. (2011) but with slight modifications as previously described in Diepenbrock et al. (2017) to account for segregation distortion across the 10 families examined herein. Only those individual-trait JL-QTL intervals for which the left and right support interval bounds uplifted continuously (among adjacent markers) and to the same chromosome from RefGen_v2 to RefGen_v4 were considered in results reporting and downstream analyses (Table S4), with the additional intervals for which the left and/or right bound did not uplift in that manner reported in Table S5.

### Genome-wide association study

First, 1,000 permutations were generated, using the residuals generated for each trait during joint-linkage analysis. From the permutation results, a significance threshold was identified for each trait corresponding to a GWAS false discovery rate of 0.05. The GWAS was run in TASSEL 4 using custom scripts (available in GitHub at time of publication). This method utilized a bootstrap resampling method in which 100 iterations of GWAS were conducted, with 80% of the RILs sampled with replacement in a given iteration. The markers tested in GWAS were the ∼26.9 million SNPs and short indels (up to 15 bp in length) of the HapMap v1 and v2 projects (Gore et al. 2009; Chia et al. 2012) that were upliftable from RefGen_v2 to RefGen_v4 coordinates. Uplifted coordinates were as determined in Diepenbrock et al. (2021), by clipping 50 markers flanking either side of a given marker (101 nucleotides in total) and aligning these to the AGP_v4 genome using Vmatch (v2.3.0; Kurtz 2019), with the following options: -d -p - complete -h1. From the alignment results, only the highest scoring and unique alignment for each marker was retained, and those markers not having a high-confidence and unique alignment were not included in the input data set for permutations and GWAS. The upliftable markers, which are available for the NAM founders, were then projected onto the NAM RILs in the 10 families under analysis in this study, using the 0.1 cM genetic linkage map, prior to permutations and GWAS. The resample model inclusion probability (RMIP; Valdar et al. 2009) was reported for each marker exhibiting a significant marker-trait association for one or more traits in GWAS, conveying in what proportion of the 100 iterations that marker was included in the final model.

### Pleiotropy

Pleiotropy was examined by fitting the final JL-QTL model for each trait using transformed BLUEs for every other trait, and then correlating the allelic effect estimates between the original trait and the other trait for every peak marker. Significance was tested with a Type I error rate of α = 0.05 using FDR-corrected *P*-values that were generated via the Benjamini-Hochberg method. These correlations were examined both within a single JL-QTL interval and at the genome-wide level, and were visualized using the *network* package in R (Butts et al. 2008; Butts et al. 2015).

### Linkage disequilibrium

Linkage disequilibrium (LD) was examined within the same HapMap v1 and v2 input data set used for GWAS. Specifically, LD was examined for markers exhibiting one or more significant associations in GWAS (hereafter, GWAS variants) as in Diepenbrock et al. (2021), by calculating pairwise correlations between GWAS variants and markers within 250 kb of each variant. A null distribution was generated by calculating the same pairwise correlations for 50,000 randomly selected variants with markers within 250 kb of the respective randomly selected variant.

### Modeling of relationships between grain carotenoid traits and kernel color

#### Linear Model

Linear models were produced using the lm() function from the lme4 package in R (Bates et al. 2015; R Core Team 2018). Each linear model was constructed as reported in Table S6. Family was treated as a categorical variable (factor). All values for traits, including both the response (kernel color) and predictors (carotenoid traits), were transformed BLUEs. The assumptions for linear regressions were checked using the plot() function in base R. The output table comparing models was produced using the compare_performance() function in the ‘performance’ package (Lüdecke et al. 2021).

#### Random Forest

Random Forest models were produced using SciKit-Learn in Python version 3.8.5 (Pedregosa *et al*., 2011; Van Rossum & Drake Jr, 1995). The transformed BLUEs were split using train_test_split from sklearn.model_selection, such that 30% of the data was used for testing, and 70% was used for training performance. Both a linear model (LinearRegression from sklearn.linear_model) and random forest model (RandomForestRegressor from sklearn.ensemble) were fit to the split training data. The random forest used 1,000 decision trees. Accuracy of the models was calculated by using r2_score (corresponding to R^2^) from sklearn.metrics. Baseline Error is defined as the absolute value of the predicted kernel color transformed BLUE minus the known test value of the kernel color transformed BLUE. Percent Accuracy is defined as 100 times the Baseline Error, divided by the test-set values of kernel color, subtracted from 100%. To add certainty that results were not affected by the randomness of the train-test-split, and to enhance reproducibility of results, the process was repeated 30 times using 30 different random seeds, for both the random forest and linear models. The model metrics were averaged over these 30 runs before being reported in Table S7. Variable Importance was extracted from the random forest model, and plotted using pyplot from matplotplib (Hunter, 2007). Feature importance was calculated over all 1000 of the decision trees, based on the average decrease in accumulated impurity, using the “feature_importances_” aspect of the random forest output.

### Data availability

Data and scripts will be made publicly available at time of publication as supplemental material, as hosted on the journal website and on GitHub, respectively.

## RESULTS

The carotenoid and kernel color traits analyzed in this study exhibited natural variation in the 10 families examined herein, with moderate to high line-mean heritabilities (Table 1). A total of 67 individual-trait JL-QTL support intervals were identified for these nine traits (Table 2). Physically overlapping intervals were combined into a single common support interval, resulting in 29 unique common support intervals. A total of 192 significant marker-trait associations (having a resampling model inclusion probability (RMIP) ≥ 5) were detected in GWAS that were contained within one of these common support intervals, with nine markers detected for multiple traits (Table S8). It was determined from the examination of linkage disequilibrium in the 10 families (Figure S1) that 250 kb on either side of a GWAS variant was a reasonable search space for gene identification. Nine genes were identified in this study, in accordance with the following criteria: residing within the search space for at least one of the 192 marker-trait associations; also residing within individual-trait JL-QTL intervals for one or more of the same traits; and having *a priori* evidence of involvement in biosynthesis of isopentenyl pyrophosphate (IPP) or biosynthesis and/or retention of carotenoids based on prior studies in plant systems (Table S9).

**Table 1.**
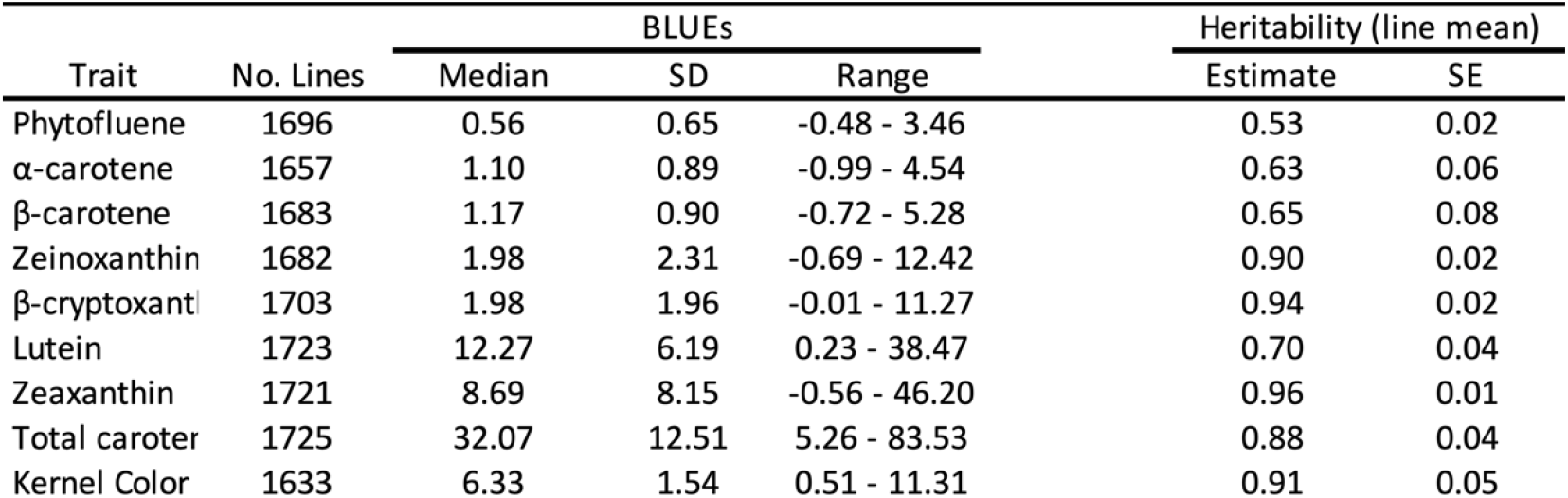
Summary statistics for untransformed Best Linear Unbiased Estimators (BLUEs) and estimates of line-mean heritability for carotenoid and kernel color traits, in the 10 U.S. maize NAM families examined in this study.

**Table 2.** Summary of joint-linkage analysis and GWAS results for carotenoid and kernel color traits in the 10 U.S. maize NAM families examined in this study.

| Trait | Number of JL-QTL | Median size (SDa ) of $\alpha = 0.01$ JL-QTL support interval (Mb) | Number of JL-QTL intervals containing a priori genes | Number of GWAS-associated variants in JL-QTL intervals | Average (Median) Percent Resampling model inclusion probability (RMIP) | Max Percent Resampling model inclusion probability (RMIP) |
| --- | --- | --- | --- | --- | --- | --- |
| Phytofluene | 9 | 11.77 (53.75) | 4 | 22 | 10.42 (7) | 46 |
| α-carotene | 4 | 9.46 (55.10) | 2 | 11 | 9.26 (7) | 57 |
| β-carotene | 4 | 6.05 (4.30) | 2 | 9 | 10.69 (8) | 47 |
| Zeinoxanthin | 7 | 5.97 (49.79) | 5 | 19 | 11.46 (8.5) | 57 |
| β-cryptoxanthin | 11 | 4.95 (4.97) | 8 | 19 | 13.61 (8.5) | 63 |
| Lutein | 8 | 6.66 (53.60) | 5 | 24 | 11.67 (7) | 64 |
| Zeaxanthin | 11 | 4.95 (49.75) | 9 | 25 | 11.62 (9) | 50 |
| Total carotenoids | 8 | 14.65 (54.44) | 3 | 21 | 11.71 (8) | 38 |
| Kernel Color | 5 | 11.77 (58.33) | 4 | 11 | 12.27 (9) | 66 |
| JL-QTL Total | 70 | 6.66 | 42 | 161 |  |  |

Three genes encoding activities in the MEP pathway, which provides substrate for carotenoid biosynthesis, were identified in this study (Figure 1, Table 3). Specifically, two homologs were identified that encode 1-deoxy-D-xylulose 5-phosphate synthase (DXS), which catalyzes the first and committed step of the MEP pathway. *dxs2* was identified with PVEs of 1.0 to 5.2% for eight of the nine traits analyzed herein, including 2.4% for kernel color (Table 3). *dxs3* was identified with PVEs of 2.2 to 3.3% for zeinoxanthin, β-cryptoxanthin, and zeaxanthin (Table 3). *mecs1* encodes methylerythritol cyclodiphosphate synthase (MECS), which catalyzes the fifth step in the MEP pathway. *mecs1* was identified with PVEs of 0.8 to 3.6% for five traits (Table 3).

**Table 3.** Percent phenotypic variation explained for each carotenoid and kernel color trait by JL-QTL resolved to each of the nine identified genes. PVEs are color-coded by the significance of the PVE, with a darker blue corresponding to a higher PVE. Trait abbreviations: phytofluene (PHYF), α-carotene (ACAR), β-carotene (BCAR), zeinoxanthin (ZEI), β-cryptoxanthin (BCRY), lutein (LUT), zeaxanthin (ZEA), total carotenoids (TOTCAR), kernel color (KCOL).

| Common Support Interval ID | RefGen_v4 ID | Annotated Gene Function | Percent phenotypic variation explained for each trait |  |  |  |  |  |  |  |  |
| --- | --- | --- | --- | --- | --- | --- | --- | --- | --- | --- | --- |
|  |  |  | PHYF | ACAR | BCAR | ZEI | BCRY | LUT | ZEA | TOT CAR | KCOL |
| 15 | Zm00001d019060 | deoxy xylulose synthase 2 (*dxs2*) | 5.2 | 2.5 | 2.9 | 3.1 | 1.9 | 1.0 |  | 3.9 | 2.4 |
| 23 | Zm00001d045383 | deoxy xylulose synthase 3 (*dxs3*) |  |  |  | 2.2 | 3.3 |  | 2.3 |  |  |
| 10 | Zm00001d051458 | methyl erythritol cyclodiphosphate synthase (*mecs1*) | 3.6 | 2.3 |  |  |  | 0.9 | 0.8 | 1.8 |  |
| 14 | Zm00001d036345 | phytoene synthase 1 (*psy1*) | 12.2 |  |  |  | 4.7 | 5.4 | 7.3 | 21.4 | 12.5 |
| 21 | Zm00001d011210 | lycopene epsilon cyclase (*lcyE*) |  | 25.1 | 15.3 | 44.0 | 29.7 | 53.2 | 29.0 |  | 17.5 |
| 3 | Zm00001d029822 | epsilon ring hydroxylase (*lut1*) |  |  |  | 4.5 |  | 2.7 |  |  |  |
| 26 | Zm00001d048469 | beta carotene hydroxylase 5 (*crtRB5*) |  |  |  |  | 1.6 |  | 1.8 |  |  |
| 6 | Zm00001d003512/<br>Zm00001d003513<sup>†</sup> | zeaxanthin epoxidase 1 (*zep1*) | 1.7 |  |  |  | 2.2 |  | 20.3 | 7.1 |  |
| 24 | Zm00001d048373 | Whitecap 1 (*wc1*) [carotenoid cleavage dioxygenase 1 (*ccd1*)] |  |  |  |  |  | 5.6 |  | 6.9 |  |

**Figure 1.**
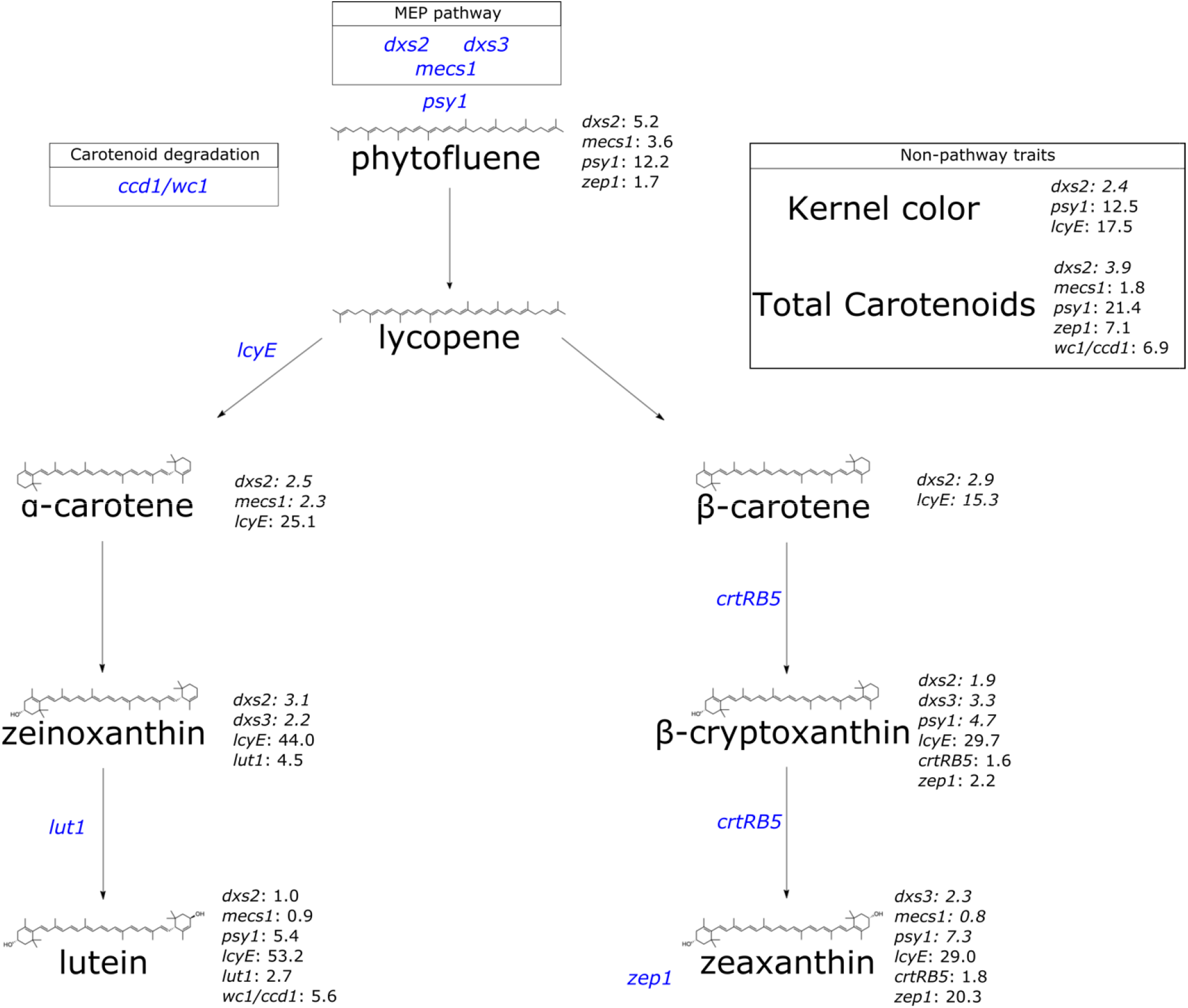
A simplified depiction of the carotenoid biosynthesis pathway, showing each carotenoid compound analyzed herein and its structure, annotated with the corresponding genes identified. Genes of interest are followed by the PVE for that gene at the corresponding trait. Gene names, listed in blue, represent the gene of interest relative to its placement in the pathway. Gene abbreviations are as follows: 1-deoxy-D-xylulose 5-phosphate synthase (*dxs2* and *dxs3*); 2-C-methyl-D-erythritol 2,4-cyclodiphosphate synthase (*mecs1*); zeaxanthin epoxidase (*zep1*); phytoene synthase (*psy1*); lycopene epsilon-cyclase (*lcyE*); Cytochrome P450 superfamily protein (*lut1*); β-carotene 3-hydroxylase (*crtRB5*); White cap1/carotenoid cleavage dioxygenase1 (*wc1*/*ccd1*).

As the committed step in carotenoid biosynthesis, phytoene synthase *(psy1)* catalyzes the synthesis of phytoene from the condensation of two 20-carbon geranylgeranyl diphosphate (GGDP) molecules. A null allele at *psy1* conditions negligible carotenoid levels in the maize endosperm (Buckner et al. 1996; Li et al. 2008). In this study, *psy1* was identified for five traits: β-cryptoxanthin (4.7% PVE), lutein (5.4%), zeaxanthin (7.3%), total carotenoids (21.4%), and kernel color (12.5%) (Figure 1, Table 3). The branchpoint in the carotenoid pathway occurs with cyclization of the ε-ring of lycopene by *lycopene epsilon cyclase (lcyE)*, which is the committed step in α-carotene biosynthesis (Cunningham et al. 1996; Bai et al. 2009; Cazzonelli and Pogson 2010). *lcyE* had the largest PVEs (15.3% to 53.2%) observed for seven of the nine traits analyzed in this study, and affected kernel color (with PVE of 17.5%) but not total carotenoids.

Within the α-branch of the carotenoid pathway, *lut1* encodes CYP97C, an ε-ring hydroxylase that catalyzes the conversion of α-carotene to zeinoxanthin and further hydroxylation of zeinoxanthin to yield lutein (Tian et al. 2004; Quinlan et al. 2012; Owens et al. 2014; Diepenbrock et al. 2021). *lut1* was identified in this study for zeinoxanthin and lutein, with PVEs of 4.5% and 2.7%, respectively (Figure 1, Table 3). Within the β-branch of the carotenoid pathway, β-carotene hydroxylase (CRTRB) preferentially converts β-carotene to β-cryptoxanthin and then zeaxanthin (Diepenbrock et al. 2021). *crtRB5* (also known as *hyd5*) was identified with PVEs of 1.6% for β-cryptoxanthin and 1.8% for zeaxanthin. Zeaxanthin epoxidase (encoded by *zep1*) converts zeaxanthin to violaxanthin, with antheraxanthin as an intermediate. *zep1* was identified with PVEs of 20.3% for zeaxanthin, 2.2% for β-cryptoxanthin, and 1.7% for phytofluene. Finally, the Whitecap locus represents a macrotransposon insertion containing some number of tandem copies of *ccd1*, which encodes a carotenoid cleavage deoxygenase (Tan et al. 2017). The *whitecap1* locus (QTL24) was identified with PVEs of 5.5% for lutein and 6.9% for total carotenoids, respectively. Overall, three of the identified genes were detected for kernel color: *lcyE* (17.5% PVE), *psy1* (12.5%), and *dxs2* (2.4%).

Pairwise correlations between untransformed BLUEs for each trait were generally moderately to strongly positive for compounds within the same pathway branch, and near-zero to negative across branches (Pearson’s *r*; Figure 2). Kernel color and total carotenoids exhibited a correlation of 0.69. Both kernel color and total carotenoids clustered with higher concentrations of β-branch carotenoids, with kernel color (and total carotenoids) specifically showing correlations of 0.76 (and 0.71) with zeaxanthin, 0.66 (and 0.54) with β-cryptoxanthin, and 0.53 (and 0.46) with β-carotene. While lutein and total carotenoids exhibited a positive correlation of 0.49, the correlation between lutein and kernel color was negligible at 0.03. Zeinoxanthin, α- carotene, and phytofluene showed only small correlations with total carotenoids, and negligible correlations with kernel color.

**Figure 2.**
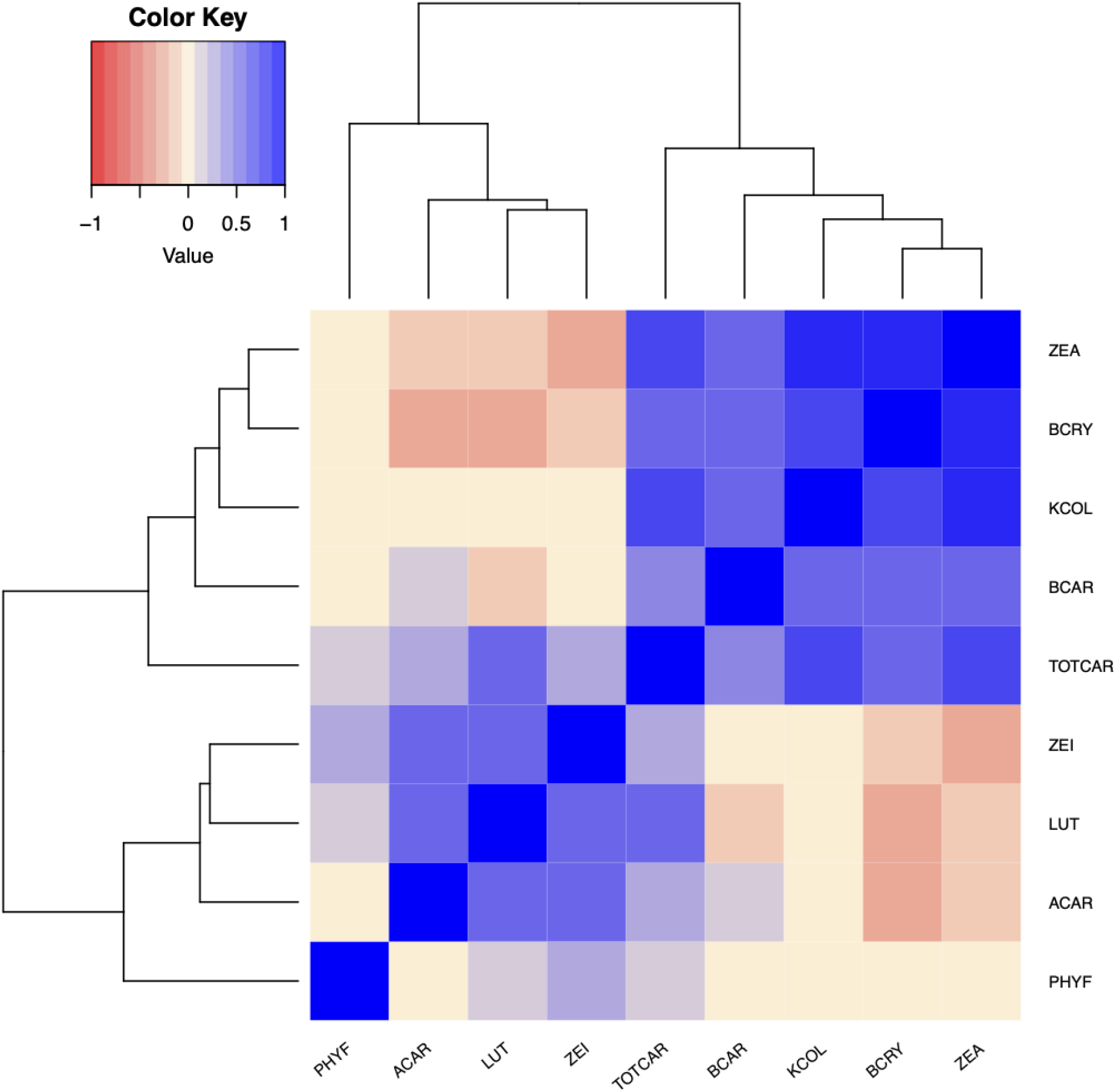
Heat map depicting correlations between untransformed BLUEs for carotenoid and kernel color traits in the 10 U.S. maize NAM families examined in this study. Trait abbreviations: phytofluene (PHYF), α-carotene (ACAR), lutein (LUT), zeinoxanthin (ZEI), total carotenoids (TOTCAR), β-carotene (BCAR), kernel color (KCOL), β-cryptoxanthin (BCRY), zeaxanthin (ZEA).

In examinations of pleiotropy within each common support interval, significant positive and/or negative pleiotropy (α = 0.05) was observed for kernel color and carotenoid traits at certain of the identified genes (Figure 3). *dxs2* exhibited positive pleiotropy for kernel color and each of β-cryptoxanthin, lutein, zeinoxanthin, and total carotenoids. *psy1* exhibited positive pleiotropy for all pairwise combinations of kernel color, zeaxanthin, and total carotenoids. *lcyE* exhibited positive pleiotropy between kernel color and each of β-carotene, β-cryptoxanthin, and zeaxanthin, and negative pleiotropy between kernel color and each of α-carotene, zeinoxanthin, and lutein. *lcyE* also exhibited negative pleiotropy for compounds across pathway branches, and for each of β-carotene and β-cryptoxanthin with zeaxanthin, which is produced downstream of these two compounds within the β-branch (Figure S2).

**Figure 3.**
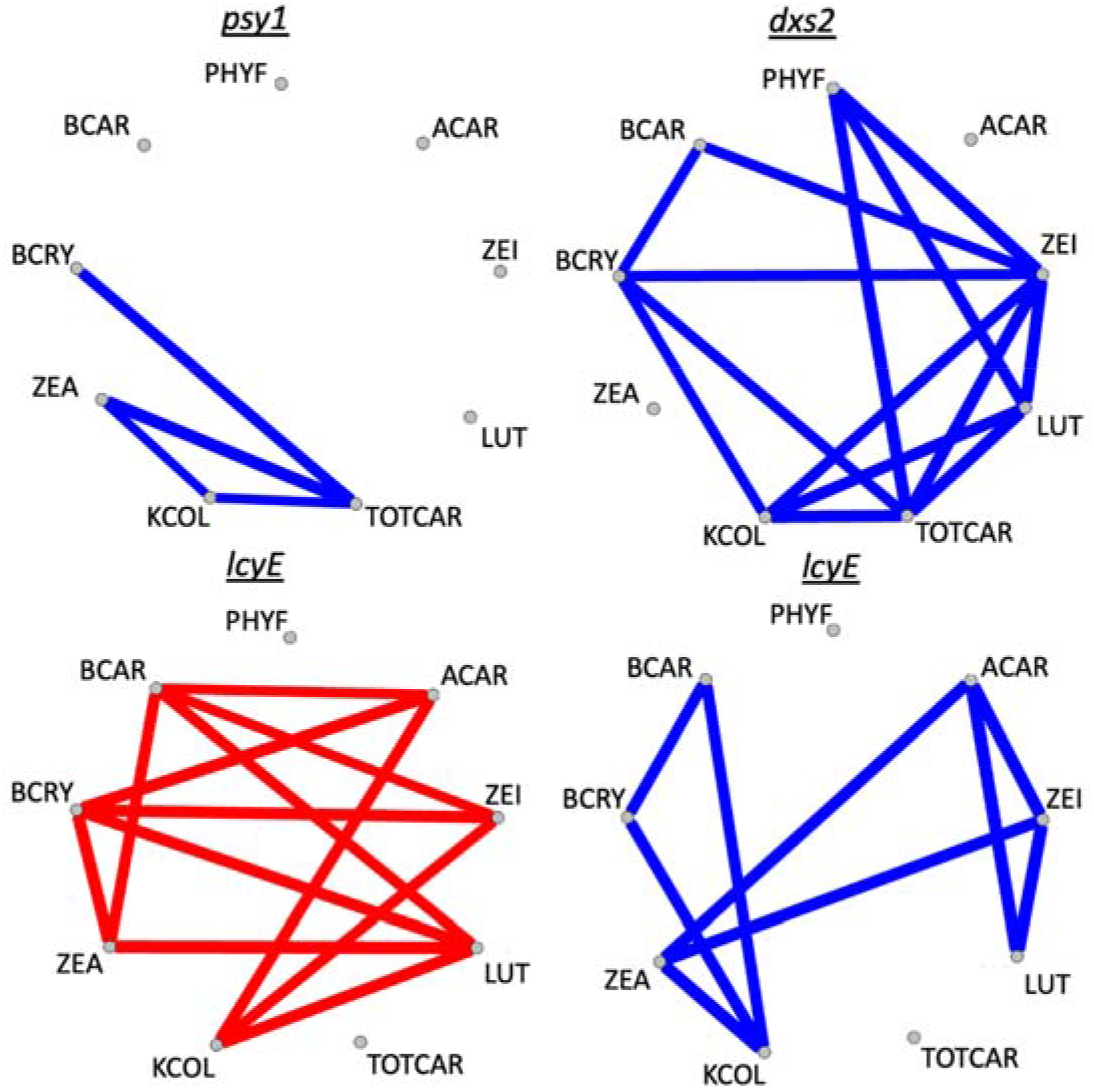
Significant pleiotropy (α = 0.05) observed for JL-QTL resolved to individual genes, between kernel color and one or more carotenoid traits in this study. Blue lines indicate positive pleiotropy, whereas red lines indicate negative pleiotropy. Trait names are as written, unless abbreviated as follows: zeinoxanthin (zeino), total carotenoids (total_carot), kernel color (traitKC), β-cryptoxanthin (b_cryp).

In modeling of kernel color as the response variable with grain carotenoid traits as predictors, the top-performing linear model was a full model that included all eight carotenoid traits analyzed in this study and a family term (the latter as a categorical variable) (R^2^= 0.727; Table S6). Models that included this family term, which contains a unique identifier (or factor level) for each of the 10 families included in this study, consistently performed better than those with the same grain carotenoid traits as predictors that omitted the family term (Table S6). The second-best performing model contained only lutein, zeaxanthin, total carotenoids, and the family term (R^2^= 0.723; Table S6). The random forest performed almost identically to a linear model fit to the same training and prediction sets (R^2^ = 0.713 and 0.719, respectively, averaged across 30 iterations; Figure S3, Table S7). The feature with highest importance in the random forest model was zeaxanthin (0.58), followed by total carotenoids (0.12) and the other two β- branch carotenoids (each at 0.05). Together, these results suggest that zeaxanthin and total carotenoids are among the top predictors of kernel color.

## DISCUSSION

This study identified QTL and marker-trait associations for eight carotenoid traits and visually scored kernel color in 10 families of the U.S. maize NAM populations, and resolved nine of these QTL to individual genes. While most of the identified genes have been previously noted as key players in carotenoid accumulation and/or kernel color (Chandler et al. 2013; Owens et al. 2014; Owens et al. 2019; Diepenbrock et al. 2021), noteworthy patterns of relevance to breeding efforts for carotenoids and/or kernel color, particularly in yellow to orange-grain maize populations, emerge from this examination of both trait sets in the same quantitative genetics-enabling experimental framework.

Within the precursor pathway, DXS has been found to be a rate-limiting activity through genetic engineering and overexpression studies in Arabidopsis and *E. coli* (Harker and Bramley 1999; Estevez et al. 2001). While *dxs2* has appeared to be the major controller in maize—with PVEs of 3.5 to 11.3% for *dxs2* and 1.2 to 4.1% for *dxs3* in the 25 NAM families (Diepenbrock et al. 2021)—these ranges were more similar in the present study: 1.0 to 5.2% for *dxs2* and 2.2 to 3.3% for *dxs3*. This could be related to *dxs2* having been found to be nearly fixed among yellow inbreds as examined in Fang et al. (2020). Nonetheless, the identification of *dxs2* and *dxs3* for carotenoid traits in both the 10 and 25 U.S. maize NAM families, and for kernel color in the 10 families as well, suggests that examination of and selection at *dxs2* and/or *dxs3* could bring about further gains. *mecs1* (encoding 2-C-methyl-D-erythritol 2,4-cyclodiphosphate synthase, or MECS) was also identified in this study for five carotenoid traits. MECS catalyzes the fifth step in the MEP pathway, which provides precursors for biosynthesis of carotenoids, tocochromanols (vitamin E-related compounds, including tocopherols), and other plastidic isoprenoids. Markers proximal to *mecs2*—another homolog encoding MECS in maize—were significantly associated with maize grain tocopherol traits in a pathway-level analysis (i.e., only testing variants proximal to *a priori* candidate genes, rather than genome-wide variants, in association analyses) (Lipka et al. 2013). Markers proximal to a gene encoding this enzymatic step were also significantly associated with grain zeaxanthin concentrations in a GWAS and pathway-level analysis in sorghum (Cruet-Burgos et al. 2020). In sum, these results indicate that DXS and MECS, potentially among other control points in the MEP pathway, could be of relevance in nutritional breeding efforts.

*Phytoene synthase 1* (*psy1*) encodes the first and committed step of carotenoid biosynthesis, and is a major controller of natural variation in grain endosperm carotenoid levels (Zhu et al. 2008; Fu et al. 2013; Diepenbrock et al. 2021). In this study, *psy1* had the largest PVE for total carotenoids and the second-largest PVE for kernel color, zeaxanthin, lutein, and β- cryptoxanthin. This finding suggests that *psy1* will continue to be an important target for selection, even among populations with yellow to orange grain (i.e., already possessing a functional allele of *psy1*). *lcyE*, which encodes an enzyme acting at the main branchpoint of the carotenoid pathway, is a major point of genetic control for levels of compounds residing in the α- or β-pathway branches (Harjes et al. 2008). *lcyE* exhibited the largest PVEs observed for seven of the nine traits, including kernel color. *lcyE* was notably not detected for TOTCAR. In combination with pleiotropy results (Figure 3), this suggests that its effect on kernel color could largely be related to its effect on flux of substrate into the two pathway branches. Within the α- branch, *lut1* was identified for zeinoxanthin (an intermediate in the reactions catalyzed by the encoded enzyme) and lutein (the product of that enzyme), which is consistent with previous findings (Diepenbrock et al. 2021; Owens et al. 2014). *lut1* was also found to exhibit negative pleiotropy between zeinoxanthin and lutein (Figure S2). *lut1* was the identified gene that was most specific to α-branch compounds, without significantly affecting β-branch carotenoids, total carotenoids, or kernel color. Within the β-branch, *crtRB5* (encoding β-carotene hydroxylase) was identified for β-cryptoxanthin and zeaxanthin. While *crtRB1* has been the homolog more frequently identified and characterized in previous studies, and with larger effect (Yan et al. 2010; Owens et al. 2014; Suwarno et al. 2015; Diepenbrock et al. 2021), *crtRB1* was not identified herein, which could be due to the populations and allele frequencies involved in this study. Further studies will need to examine the role of *crtRB1* with regards to the relationship between kernel color and grain carotenoid traits. Finally, *zep1* was identified in the present study for three traits, explaining the greatest phenotypic variation for zeaxanthin (20.3% PVE). Examination of allele and haplotype frequencies and allele mining for the nine genes identified in this study will be an important next step in simultaneously optimizing kernel color and carotenoid traits in breeding populations.

The overall PVE ranking for kernel color in this study was *lcyE* (17.5%), *psy1* (12.5%), and *dxs2* (2.4%) (Table 3, Figure 1). Of these, *dxs2* exhibited several positive and no negative pleiotropic relationships for the traits in this study (Figure 3). *psy1* exhibited a smaller number of positive pleiotropic relationships, and one instance of negative pleiotropy between β- cryptoxanthin and zeaxanthin. *lcyE* was found to exhibit negative pleiotropy between kernel color and α-branch carotenoids (and positive between kernel color and β-branch carotenoids). Thus, if directing flux towards the β-branch of the carotenoid pathway is desirable, selecting on *lcyE* for kernel color could be advantageous. Alternatively, if increased concentrations of certain α-branch compounds, such as lutein, are also desired, it appears that *dxs2* and *psy1* could be used to improve kernel color (and lutein) without imposing such tradeoffs. In that case, the allele to be selected upon at *lcyE* could then be determined based primarily on the balance of α- and β- branch compounds that is optimal for human nutrition in the desired use case. Notably, negative pleiotropy was also observed within the β-branch for *lcyE*, which was not observed in the 25 NAM families (Diepenbrock et al. 2021). It will be important to monitor whether selection at *lcyE* within yellow-to orange-grain breeding populations results in tradeoffs among β-branch compounds—namely given that *lcyE* has been one of two genes targeted in marker-assisted selection, and given the large PVEs exhibited by *lcyE* for β-branch carotenoids (among other priority traits). Such tradeoffs could potentially be mitigated through selection at other genes exhibiting large PVEs for β-branch carotenoids (Table 3).

The three genes identified for kernel color in the present study—*lcyE, psy1*, and *dxs2*— were also identified in Owens et al. (2019). That study examined quantitative components of kernel color (as specified in the three-dimensional CIELAB color system, which uses pairs of color opponents) in 1,651 inbreds of the Ames national maize collection via handheld colorimeter, and identified *lcyE* for hue angle (representative of perceived color), *psy1* for *a*\* (greenness to redness), and *dxs2* for both *a*\* and hue angle. The fourth gene identified at the genome-wide level in Owens et al. (2019), for *a*\*, was *zep1*, which was identified for three traits herein but not for kernel color. While there were GWAS variants for kernel color for which *zep1* was in the ±250 kb search space (Table S8), an individual-trait JL-QTL interval was not identified for kernel color in the vicinity of *zep1* in RefGen_v4 coordinates (Table S5). Intervals were identified for kernel color in RefGen_v2 coordinates that contained *zep1* (PVE 3.0%) as well as *dxs3* (PVE 4.3%; Table 3), but one support interval bound for each of these intervals uplifted to a contig rather than to the same chromosome (Table S5), such that those intervals were not included in downstream analyses. Future studies involving these two genes could likely still benefit from continued monitoring of potential impacts on kernel color.

The Chandler et al. (2013) study that initially identified QTL for the visually scored kernel color trait examined herein, using 1,104 genetic markers and without having grain carotenoid data in the 10 families, also identified *lcyE* (PVE of 38.3%) and *psy1* (20.7%) as well as *zep1* (6.6%) and *wc1*/*ccd1* (8.7%). A JL-QTL support interval was detected in the present study for kernel color (PVE 2.9%) which contained the *ccd1-r* progenitor locus. However, there were not significant marker-trait associations for kernel color within that interval (rather, on either side), such that the interval was not resolved to the gene level. The macrotransposon insertion site containing a tandem number of copies of *ccd1*, located 1.9 Mb upstream of the *ccd1-r* progenitor site (Tan et al. 2017), was identified in this study with moderate PVEs for lutein and total carotenoids. Given the multiple evident signals and somewhat dispersed nature thereof, further examination of *Whitecap* and the *ccd1-r* progenitor region is merited in breeding populations, including those having yellow to orange grain. Colorimetric examinations of kernel color could additionally be helpful in future efforts, for purposes of distinguishing the effects of genomics-assisted breeding efforts for orange, carotenoid-dense maize on multiple quantitative components of kernel color—namely *a*\* and hue angle, for which genetic associations were detected in Owens et al. (2019), as well as other traits quantified in the CIELAB color system, such as chroma (i.e., color saturation or intensity). An examination of zeaxanthin-biofortified sweet corn found hue angle to decrease from 90° (noted as yellow) to 75° (noted as yellow-orange) with increasing concentrations of β-branch carotenoids (O’Hare et al. 2015).

Pigment color is a signature feature of different carotenoids, and is directly mediated by desaturation reactions occurring throughout the biosynthetic pathway (Bartley and Scolnik 1995; Khoo et al. 2011; Meléndez-Martinez et al. 2007). Namely, increasing the number of conjugated double bonds affects the chromophore comprising the carotenoid backbone and determines the exact wavelengths of light that it is able to absorb and emit (Bartley and Scolnik 1995; Meléndez-Martínez et al. 2007; Saini et al. 2015). Generally, carotenoids that carry more conjugated double bonds (such as lycopene) maximally absorb longer wavelengths of light and are deeper in hue than other carotenoids such as phytoene and phytofluene, which have only three and five conjugated double bonds, respectively, and are effectively colorless. Indeed, zeaxanthin contains 11 conjugated double bonds and appears as a deeper orange color relative to lutein, which has 10 conjugated double bonds and emits a more yellow color (Bartley and Scolnik 1995; Khoo et al. 2011; Meléndez-Martínez et al. 2007). This trend is consistent for the other β- and α-branch compounds examined herein: β-carotene and β-cryptoxanthin have a carbon backbone of 11 conjugated double bonds and appear more yellow-orange in hue compared to α-carotene and zeinoxanthin, which contain only 10 conjugated double bonds and are paler yellow carotenoids (Khoo et al. 2011; Meléndez-Martínez et al. 2007).

It is thus not surprising that kernel color was more strongly correlated with β-branch carotenoids in these populations with yellow to orange grain. The results of this study indicate that zeaxanthin was the primary driver of this relationship, given its feature importance in the random forest model (0.58, compared to 0.05 for each of the other two β-branch compounds analyzed herein: β-carotene and β-cryptoxanthin; Figure S1), and its higher abundance compared to—and positive correlations with—these two compounds. Negligible correlations between kernel color and α-branch carotenoids were less expected. At least weak correlations could have been feasible, given the abundance of lutein in grain of diverse maize accessions (Owens et al. 2014) and its moderate positive correlation (*r* = 0.49) with total carotenoids as observed in the present study. Stronger relationships between kernel color and α-branch carotenoids than those observed in this study could likely be expected if white-endosperm lines were to be included in the populations under examination.

The random forest model performing almost identically to the linear model for prediction of kernel color via grain carotenoid traits suggests a linear relationship between carotenoid traits and kernel color in the populations examined herein. Zeaxanthin and total carotenoids were identified as top predictors of kernel color both in the best-performing linear models and in the random forest feature importance analyses. Sweet corn lines with higher zeaxanthin (and more generally, β-branch carotenoid) levels were also found to have a deeper orange perceived kernel color (in the form of hue angle; O’Hare et al. 2015). Together, these results suggest that breeding for zeaxanthin would also be a viable method to breed for orange kernel color in maize. This tandem improvement is aligned with breeding priorities to increase concentrations of zeaxanthin and lutein alongside provitamin A, as priority carotenoids for human health.

Finally, kernel hardness (or vitreousness, rather than opacity) is an important agronomic and processing trait, and is also a relevant consideration with regards to carotenoids and kernel color. A non-functional allele of *crtRB1—*a gene not identified herein*—*was recently found to confer kernel opacity (Wang et al. 2020). *dxs2* and *seed carotenoid deficient 1* (which encodes the enzymatic step downstream of that encoded by *mecs1*, identified herein) were detected as modifiers in that background, as they conferred altered endosperm color and kernel vitreousness (Wang et al. 2020). *zep1*, also identified herein, was additionally detected in that study as a candidate modifier for kernel vitreousness in that background. Notably, kernel hardness (which corresponds to greater transmission of light) has also been found to affect perceived kernel color (Saenz et al. 2020; Saenz et al. 2021). Linear models predicting kernel color from grain carotenoid traits were found to have R^2^ of 0.65 to 0.66 without a family term, and 0.71 to 0.73 with a family term. These results indicate that grain carotenoid traits are moderately predictive of kernel color. Kernel hardness and other physical/structural properties may explain some of the remaining ∼30% of phenotypic variation for kernel color. Examination of ground kernels (or flour) in future studies would partially remove this potential confound arising from physical/structural relationships and could be of relevance to additional use cases for maize grain. When improving grain carotenoid traits and whole-grain kernel color in tandem, including through selection at one or more of the genes identified in this study, it would be helpful to monitor kernel hardness, both to ensure maintenance of this important trait and to further improve our understanding of relationships between the abundance/composition of grain carotenoid traits and final perceived kernel color, in breeding populations and developed maize varieties.

## CONCLUSION

This study simultaneously investigated the genetic basis of grain carotenoid traits and visually scored kernel color, in the same experimental framework (field environments and populations). Nine genes were identified for these traits, which encode activities in the precursor and core carotenoid pathways as well as carotenoid degradation. The genetic and phenotypic relationships dissected herein provide additional refinement—including key genes, relevant synergies and tradeoffs, and other potential look-out points—for breeding of orange, carotenoid-dense maize, including from germplasm pools having yellow to orange grain.

## ACKNOWLEDGEMENTS

This work was funded by a U.S. Department of Energy Computational Science Graduate Fellowship under award number DE-SC0021110 (MFL) and by UC Davis start-up funds (CHD).

## AUTHOR CONTRIBUTIONS

MFL and CHD planned and designed the research. MFL, MV, MF, and CHD conducted the analyses and wrote the manuscript. MV and MF contributed equally.

